# Regional protein expression in human Alzheimer’s brain correlates with disease severity

**DOI:** 10.1101/283705

**Authors:** Jingshu Xu, Stefano Patassini, Nitin Rustogi, Isabel Riba-Garcia, Benjamin D. Hale, Alexander M Phillips, Henry Waldvogel, Robert Haines, Phil Bradbury, Adam Stevens, Richard L. M. Faull, Andrew W. Dowsey, Garth J. S. Cooper, Richard D. Unwin

**Affiliations:** Division of Cardiovascular Sciences, School of Medical Sciences, Faculty of Biology, Medicine and Health, The University of Manchester, Manchester, M13 9WL and Centre for Advanced Discovery and Experimental Therapeutics (CADET), Central Manchester University Hospitals NHS Foundation Trust, Manchester Academic Health Sciences Centre, Manchester, UK; School of Biological Sciences, and Maurice Wilkins Centre for Molecular Biodiscovery, Faculty of Science, University of Auckland, New Zealand; Department of Electrical Engineering and Electronics, University of Liverpool, Liverpool, L69 3GJ, UK; Centre for Brain Research, Faculty of Medical and Health Sciences, University of Auckland, Auckland, New Zealand; Research IT, The University of Manchester, Manchester, UK; Division of Developmental Biology & Medicine, School of Medical Sciences, Faculty of Biology, Medicine and Health, University of Manchester, Manchester, M13 9PL, UK; School of Social & Community Medicine and School of Veterinary Sciences, Faculty of Health Sciences, University of Bristol, Bristol BS8 2BN, UK

## Abstract

Alzheimer’s disease (AD) is a progressive neurodegenerative disorder that currently affects 36 million people worldwide with no effective treatment available. Development of AD follows a distinctive pattern in the brain and is poorly modelled in animals. Therefore, it is vital to widen both the spatial scope of the study of AD and prioritise the study of human brains. Here we show that functionally distinct human brain regions show varying and region-specific changes in protein expression. These changes provide novel insights into the progression of disease, novel AD-related pathways, the presence of a ‘gradient’ of protein expression change from less to more affected regions, and the presence of a ‘protective’ protein expression profile in the cerebellum. This spatial proteomics analysis provides a framework which can underpin current research and opens new avenues of interest to enhance our understanding of molecular pathophysiology of AD, provides new targets for intervention and broadens the conceptual frameworks for future AD research.

---

Alzheimer’s disease (AD) is a multifactorial neurodegenerative disorder characterized by progressive dementia^1,2^. Accumulation of Aβ peptide and microtubule-associated protein tau, which exhibits hyperphosphorylation, and oxidative modifications into so-called ‘plaques’ and ‘tangles’ are considered to be central to the pathology of AD^3^. Other prominent features of AD include early region-specific decline in glucose utilisation and mitochondrial dysfunction and consequently depleted ATP production and increased reactive oxygen species production in neurons^4^. Excitotoxicity in the AD brain arising from altered glutamatergic signalling^5^, and dysregulation in other neurotransmitters has also been documented, including abnormalities of adrenergic, serotonergic and dopaminergic neurotransmission^6^. In response to pathological stimuli associated with AD, inflammatory events mediated through both innate and cell-mediated immune mechanisms are also present^3^.

Despite an increase in research into the underlying pathology of AD over the last decade, there remains controversy around what underpins this disease process, which in turn affects the pipeline of new disease modifying agents. There remains a lack of detailed mechanistic knowledge about what happens in the human brain in AD. This is exacerbated by the fact that different brain regions develop pathology at different times in the disease process, adding a spatial element to the disease which is not captured by work in cell culture models and is often overlooked in human studies, which tend to focus on single regions. Animal models also fail to capture the full disease process, at either the behavioral or biochemical levels^7^, such that translation of both basic biological findings and/or the activity of potential disease-modifying interventions from animals into humans is relatively unsuccessful. While there have been several studies which have focused on the transcriptome in human AD, there is a wealth of evidence that suggests many protein expression changes in biological systems can occur independently of transcript-level regulation, and that studying the proteome can prove new insights on the regulation of functionally active molecules in a given biological or disease state^8^.

Mass spectrometry based proteomics has been recognised as a powerful tool with the potential to uncover detailed changes in protein expression^9^. To date, however, there are few studies of protein expression in AD carried out using human brain tissue, and those that exist typically examine a single AD affected brain region^10,11^, and use different patient cohorts and analytical methods that makes between-region comparisons difficult. Such studies also frequently use either small numbers of samples (n<4) or cohorts poorly matched for age or tissue *post-mortem* delay^10,12,13^. This study aims to overcome some of these existing limitations by providing a spatially-resolved analysis of protein expression in six regions of human control and AD-affected brain in well matched, short *post-mortem* delay tissue.

## Results

In this study, we analysed six functionally distinct regions of human *post-mortem* brain; hippocampus (HP), entorhinal cortex (ENT), cingulate gyrus (CG), sensory cortex (SCx), motor cortex (MCx), and cerebellum (CB), by mass spectrometry to gain a more comprehensive understanding of protein expression changes within the AD brain. These regions were selected to represent parts of the brain known to be heavily affected (HP, ENT, CG), lightly affected (SCx, MCx) and relatively ‘spared’ (CB) during the disease process. Donors were well matched for age and *post-mortem* delay times were short, with no significant difference between cases and control. Donor data is provided in Supplementary Table 1. Relative protein expression was determined using an isobaric tagging approach followed by 2-dimensional liquid chromatography and mass spectrometry. Peptide-level data were then analysed using a Bayesian model that infers a posterior probability distribution for the relative levels of each protein between ‘cases’ and ‘controls’ based on the underlying relative peptide levels. To promote sharing and usage of these data, we have developed a searchable web interface that hosts all of our results (www.manchester.ac.uk/dementia-proteomes-project; described in Supplementary Information), which also includes Bayesian probability distributions for each protein across all individual brains examined in this study. The complete workflow is illustrated in Figure 1. The complete processed data for each region (at protein identification FDR <1%) can be found in Supplementary Table 2. Raw mass spectral data can be accessed via PRIDE, with initial search outputs prior to Bayesian modelling available via the Open Science Framework at DOI 10.17605/OSF.IO/6BXJQ (Supplementary methods).

**Figure 1.**
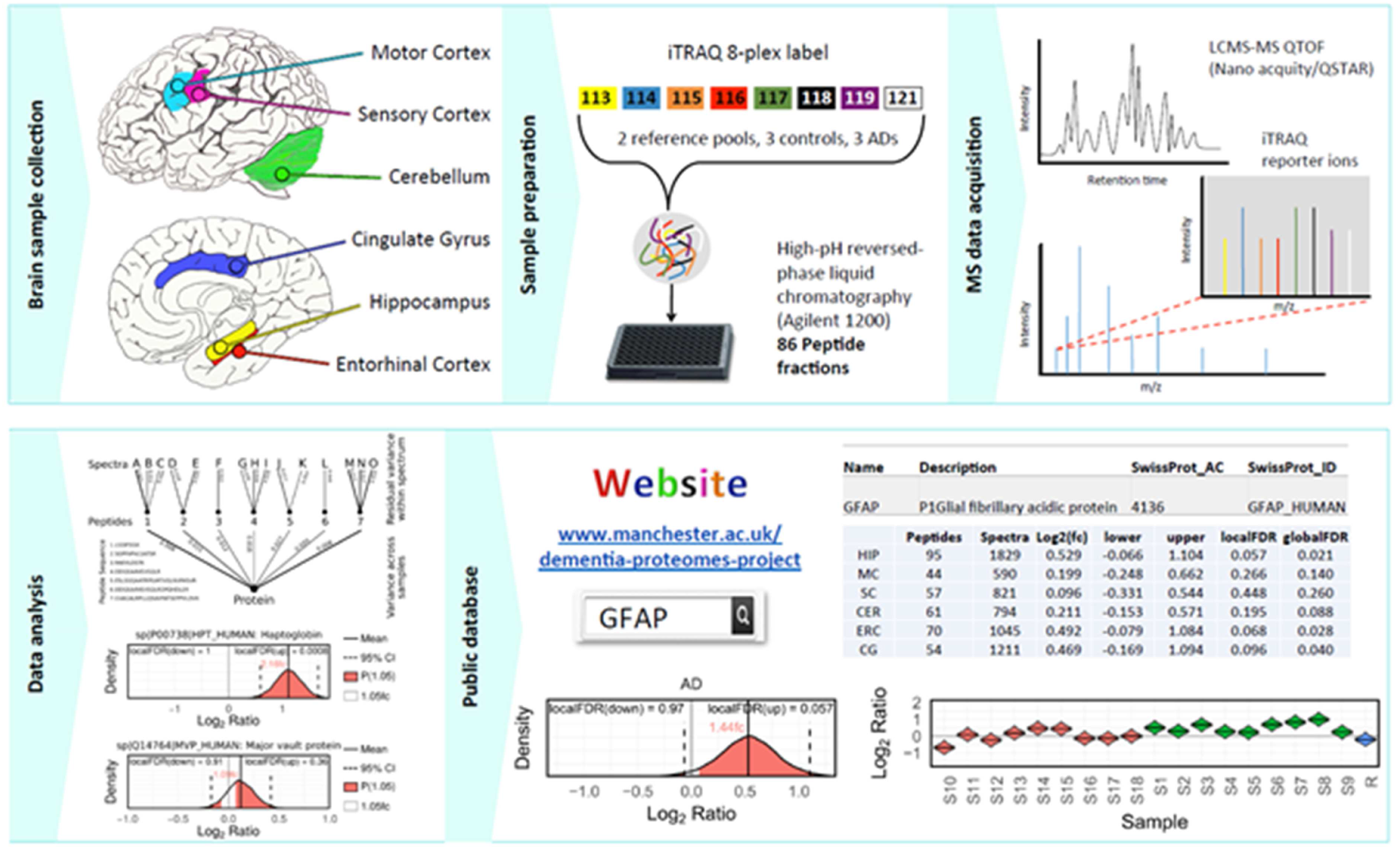
Proteomics workflow. Selected brain regions were pre-dissected prior to storage at −80°C until analysis. Each region was lysed, and protein assigned to an iTRAQ 8plex. Following digestion and labelling, samples were pooled, peptides fractionated by High-pH reverse phase chromatography and fractions analysed by standard LC-MS/MS methods. Peptides were identified and quantified based on their iTRAQ reporter area; relative protein quantification was inferred from these values using a Bayesian model. All data are deposited in a searchable online database.

Each brain region was analysed in isolation, adding strength to our comparison of protein expression changes across multiple regions, since these were identified and quantified independently. Combining all protein identifications (at 1% false discovery rate) across the six experiments yielded a total of 5,825 unique protein identifications across all regions. In our data, 990 proteins were quantified with only one or two spectra in any single region, and were subsequently omitted from our downstream cross-regional comparison in order to retain the proteins with the most precise quantification – optimisation data suggests that when the same sample is split and processed independently, >99% of proteins are defined as not being significantly different above this threshold (data not shown). However, many of these will be quantified correctly (we have previously validated expression changes based on a single spectrum, e.g. p53 in^8^), and as such these data have been included in Supplementary Table 2 and our online database. We thus quantified a total of 4,835 distinct proteins in at least one brain region, among which 3,302 proteins were common to at least three regions, and 1,899 to all six regions (Fig. 2a). These data allow us to a) define protein changes as a result of AD in any given region of the human brain being studied, and b) identify differences in how distinct brain regions are affected in AD, and by extension protein changes which occur in multiple regions of the AD brain.

**Figure 2.**
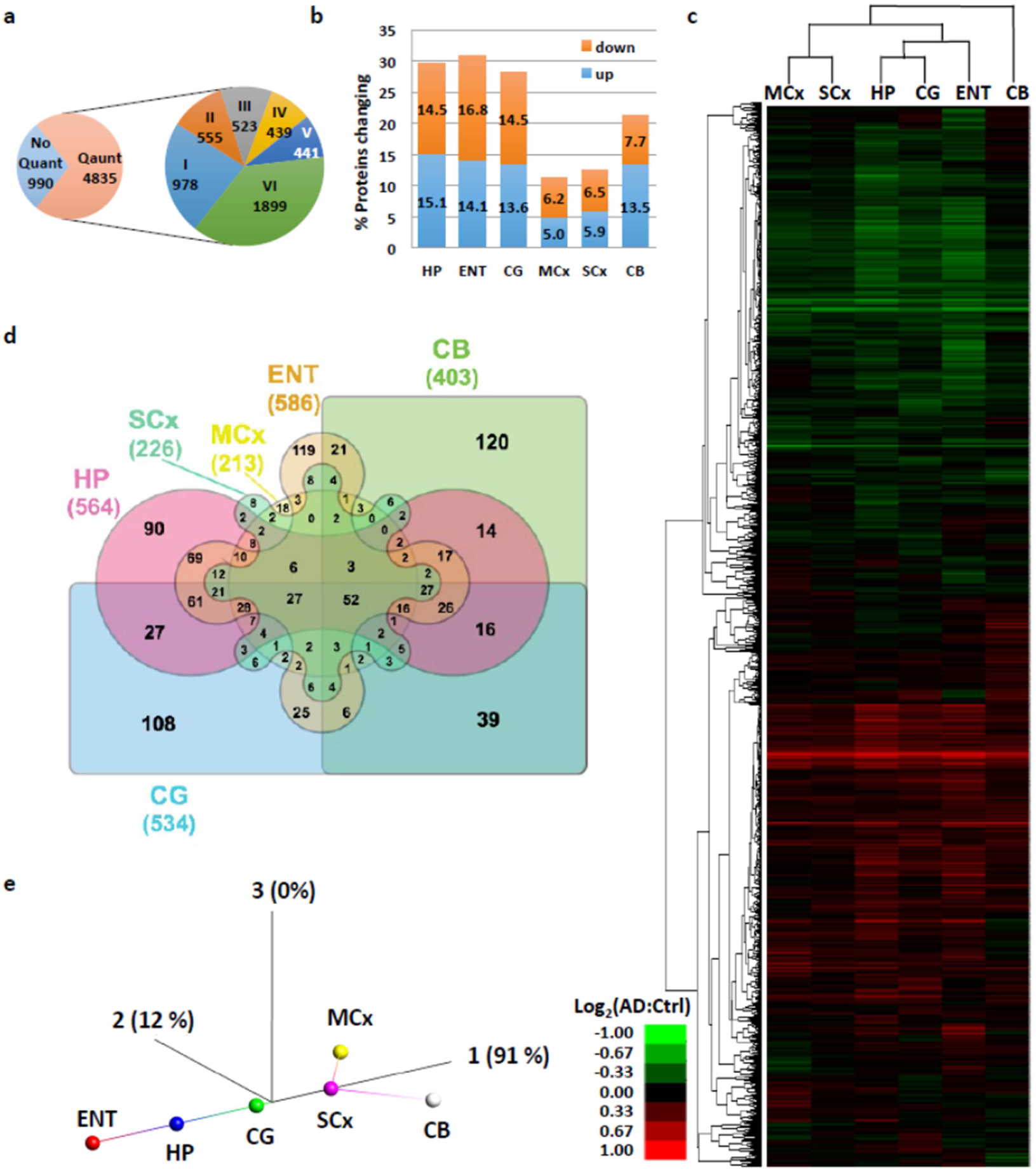
Summary of protein expression data. a) 5,825 proteins were identified, with 990 quantified with only one or two spectra and which were thus omitted from our primary comparative analysis. The remaining 4,835 proteins are classified as to whether they were quantified in six or fewer distinct regions. b) Proportion of identified, quantified proteins showing a change in expression in AD in each of the six regions under study. c) Heat map and dendrogram showing the relationship between protein expression in each region mapped using proteins present in all six regions, with three distinct ‘groups’ based on highly affected (HP, ENT, CG), moderate (MCx, SCx) and spared (CB) clearly visible. d) Edwards-Venn diagram showing the overlap of protein expression changes between brain regions, including only proteins quantified in all regions. e) Isometric mapping (Isomap) representation of protein expression data between brain regions showing a broadly linear relationship from non-affected towards affected regions, with the exception of cerebellum, which shows distinct patterns of protein expression in AD.

Comparison of the total number of proteins whose expression is altered in each region reveals, perhaps unsurprisingly, that the more severely affected areas in AD (HP, ENT, CG) show the largest number of changes in protein expression (∼30% of quantified proteins), while less affected regions (MCx, SCx) have fewer changes (11-13%). Strikingly, the CB, which many think to be pathologically ‘unaffected’, shows a substantial number of protein changes (20%; Fig. 2b). This observation accurately recapitulates data from our previous study of the metabolome on these brain samples^14^. Unsupervised hierarchical clustering of protein expression changes from all six regions demonstrates that the changes observed in CB are distinct from those seen in the affected HP, CG and ENT (Fig.2c). This is supported by an Edwards-Venn representation of the data which shows that 120/403 (29.8%) of changes in CB are not seen elsewhere (Fig.2d; Supplementary Table 3). While it has long been reported that the CB in AD can contain amyloid plaques^15^, it is considered to be relatively ‘spared’ in AD. There is a lack of neurofibrillary tangles in cerebellum^16^, and this region does not appear to develop significant neuronal loss, such that this region is often used as a control in imaging studies of the AD brain^17,18^. However, recent work by Guo *et al.* suggests a distinct pattern of cerebellar atrophy, which spreads from intrinsic connectivity networks within the cerebrum^19^, and alterations in cerebellar glucose metabolism have been reported in late stages of the disease^20,21^. Our data strongly suggest that the CB is heavily affected by AD at the molecular level, at least in late stage disease, and is so to a greater extent than other regions associated with later degeneration such as MCx or SCx, where protein changes were fewer and encompass those seen in the more severely affected regions. That the changes in CB are different from those seen elsewhere in the brain raises the possibility that, rather than being ‘spared’, the CB is affected in a different way to other brain regions and that, given it shows little pathology, these changes may reflect some level of active protection.

Hereinafter, we refer to HP, ENT and CG as the severely affected, and MCx and SCx as the less affected regions based on the number of significantly altered proteins and pathways observed within this study.

Unsupervised clustering of brain regions based on their protein expression, by performing a dimensionality reduction on these data using isomeric feature mapping (Isomap), clearly shows this hypothesized ‘evolution’ of the disease from the least affected cortical regions to the most affected, with cerebellum following a distinct pathway from the inception of disease (Figure 2e). This non-linear approach has been shown to be an improvement over the more standard PCA approach for analysis of gene and signalling networks^22^. These data also further support our previous observation that CB stands out as a single, uniquely affected brain region based on the distinctive patterns of changes found here while the other regions line up along the same vector in accordance with disease severity. Previous studies using gene co-expression networks and transcriptomics analysis have demonstrated a pattern where the molecular signatures in less-affected areas of the brain overlap with but are less marked than the grossly affected areas, and have implied that these overlapping changes represent those which occur early in AD-related neurodegeneration^23^. Our data at the protein level would support this conclusion - the less affected regions (MCx and SCx) contain very few protein changes which are not seen elsewhere, and a clustering analysis suggests that these regions are simply at an earlier stage down a similar pathway. Therefore, our data shows that by comparing more and less affected brain regions in a multi-regional approach we can observe different stages of the same disease process, enabling identification of early molecular changes, even in patients with late-stage disease.

To probe the differences in AD-related protein expression between brain regions in more mechanistic detail, we performed a pathway enrichment analysis for all differentially expressed proteins for each region. Such analyses enable us to visualise which processes are affected in the AD brain, and also whether two (or more) regions are showing dysregulation in the same pathway even if different subsets of proteins are identified as ‘changing’. These data are summarised in Figs 3a-f (and Supplementary Table 4).

**Figure 3.**
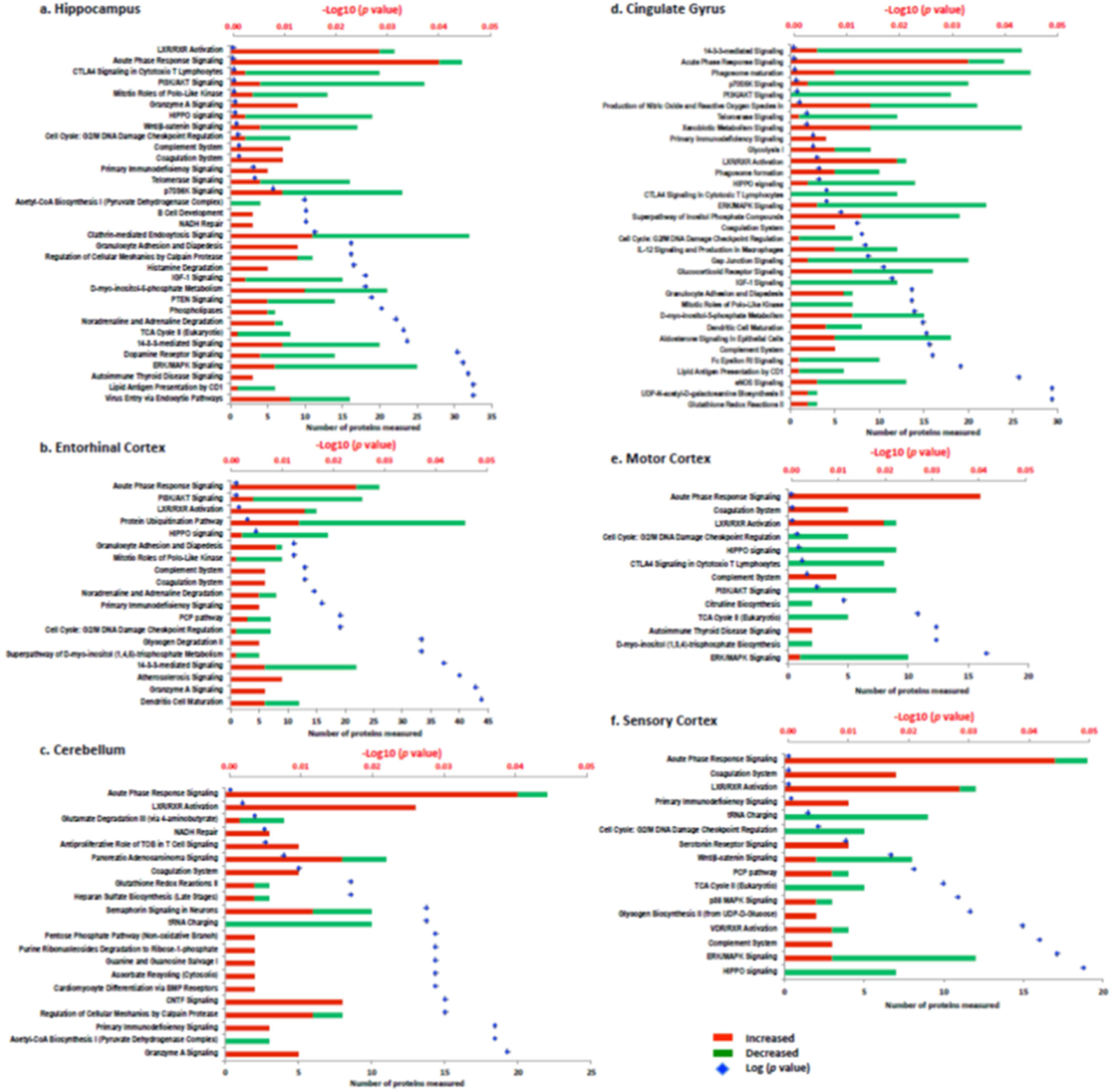
Network analysis summary. Alterations of molecular pathways in human Alzheimer’s disease brain across six distinct regions, namely a) Hippocampus, b) Entorhinal cortex, c) Cerebellum, d) Cingulate gyrus, e) Motor cortex, and f) Sensory cortex. In each plot, the numbers of increased and decreased proteins are indicated by the red/green bars, while the blue spots indicate the log_10_(p-value) for each pathway.

Reflecting the individual protein expression data, HP and CG showed the highest number of biological pathways being affected by AD. The changes in specific molecular pathways were comparable between HP, ENT, and CG. CB, on the other hand, showed altered regulation of a set of molecular pathways with limited overlap with those affected in the other five brain regions, again arguing for the presence of a distinct cellular response to disease in this region.

One of the most consistent features across all brain regions was a significant change in proteins and pathways involved with the innate immune response. In AD, aggregates of Aβ can trigger both pathogen-associated and initiate immune responses, and a persisting elevation of Aβ may elicit a chronic reaction of the innate immune system^24^. In this study, we observed strong evidence for the global activation of the innate immune response, including of the acute-phase response, the complement system (classical and alternative pathways) and the coagulation system, consistent with widespread neuroinflammation, suggesting that this may be a relatively early (prior to atrophy) event in pathogenesis. Previous studies have also implicated complement family proteins as potential AD biomarkers^25^, and GWAS studies have identified AD risk loci in a number of complement pathway genes^26-28^. It is worthy of note that these studies do not directly inform on the activation state of the complement pathway, and indeed in our study we see upgregulation of SerpinG1, which inhibits complement C4 cleavage by C1 and MASP2, as well as increased levels of C4, C3 and various regulators in AD. While it is highly likely that dysregulation of this pathway plays a role in AD, the precise nature of this role remains to be determined. Overall, HP, ENT and CG showed substantive evidence for a broader spectrum of changes in immune responses compared to MCx, SCx and CB. These included specific cellular pathways including granulocyte adhesion and dendritic cell maturation (Fig. 3a–f, Supplementary Data Table 4 and 5), implying that the innate immune system becomes activated early, and that the adaptive immune response plays a role later in the disease process. However the interplay between these two systems is complex and it is yet to be determined if these changes are a cause, or a consequence of other aspects of AD pathogenesis^29^.

This pathway-level analysis also identified signaling pathways involved in apoptosis and cell cycle regulation as being widely dysregulated in severely affected regions of AD brain, including the HIPPO, ERK/MAPK, PI3K/AKT, and Wnt/β-catenin pathways (Fig.3a, b, d), all known to be critically involved in regulation of apoptosis and the cell cycle. Reduced abundance of proteins involved in Polo-Like Kinase signaling and G2/M DNA Damage Checkpoint Regulation are likely a cause of impaired cell cycle regulation, marking these pathways out as potentially key contributors to neuronal cell death in AD. Strikingly, less affected regions SCx and MCx do not show large changes in these pathways (Fig.3e, f), reflecting reduced levels of apoptosis seen in these areas. The only exceptions are the G2/M checkpoint and the Hippo pathway, whose members are significantly decreased in these regions, suggesting that inactivation of this key developmental pathway, possibly via the observed upregulation of CD44^30^, or altered regulation of associated proteins such as the synaptic scaffolding proteins DLG2, DLG3, and DLG4, all of which are downregulated, is an early event in AD development. In CB, only granzyme A signaling was identified as an apoptosis-related pathway, indicative of fewer cell death signals in this region.

We also observed both global and regional metabolic impairments in the AD brain. Defects in brain metabolism and energetics are central to the pathogenesis of AD as evidence by epidemiological, neuropathological, and functional neuroimaging studies^31^. The AD brain characteristically exhibits defective cerebral perfusion^32^ and glucose uptake^33^, which is believed to underlie hypometabolism and cognitive decline^34^. Alterations in pathways of monosaccharide/glucose metabolism are highly significant in severely affected brain regions and CB (Fig.3a – f, Supplementary Data Table 4), consistent with our previous finding of elevated free glucose levels in AD brain^21^. TCA enzyme abundance was generally decreased in all regions of AD brain, going some way to explaining the previously observed shift from primarily aerobic glycolysis (i.e. glycolysis followed by complete oxidation in mitochondria) to the ketogenic/fatty acid β-oxidation pathway, with impaired mitochondrial bioenergetics^35^. Severely affected brain regions also showed substantial alterations in signals related to altered regulation of neurotransmitters/hormones (noradrenaline/adrenaline, dopamine, and aldosterone) that were not observed in less affected regions. While this might suggest that altered neurotransmitter biology is a late or downstream process in pathogenesis, it is notable that the enzymes in a key upstream pathway of neurotransmitter production which results in the production of tetrahydrobiopterin (BH4), a precursor of dopamine, noradrenaline and serotonin, is significantly upregulated in all regions studied. Previous work has suggested a decrease in BH4 levels in AD brain^36^ and the observations at the protein level may reflect a feedback loop where the cell is responding to decreased BH4. The presence of this dysregulation early in disease suggests it is a target which deserves closer attention.

While comparison of affected regions yields a range of interesting and novel observations about the molecular underpinning of AD, the presence of a large number of changes in ‘unaffected’ cerebellum provides a surprising finding, even more so when one observes that these changes are distinct from those manifest elsewhere. To investigate this population of protein changes further, we analysed proteins uniquely affected in CB using both DAVID and STRING. These analyses supported our earlier global pathway analysis in demonstrating that CB additionally showed alteration in Semaphorin and ciliary neurotrophic factor (CNTF) pathway members which play important roles in neuronal survival and neurodevelopment/neuronal regeneration (Fig.3c and Fig.4a, b). SEMA7A, shown here to be upregulated in CB of AD brains, is known to be involved in repair of the glial scar following spinal cord injury and to play a role in the development of multiple sclerosis, but has not previously been linked to the disease process in AD^37^. CB also showed a significant reduction in levels of both nuclear and mitochondrial aminoacyl-tRNA synthetases. In CB, significantly depleted aminoacyl tRNA synthetases, including those encoded in the mitochondrial genome as well as those from the nuclear genome (Fig.3c and Supplementary Data Table 3), could disrupt translational fidelity, leading to accumulation of misfolded proteins^38^. However, these proteins are multifunctional. For example, Ishimura *et al.* have shown that misregulated tRNA processing can lead to neurodegeneration^39^, and tRNA synthetases have also been shown to be mediators of inflammation^40^ thus downregulating these proteins may confer some level of protection. This finding could also provide a supportive mechanism for the hypothesis that ribosomal dysfunction is an early event in AD^41^. Taken together with its known roles in inflammation and signaling, and in several other neurodegenerative disorders^42^, our data suggest that the role of tRNA synthetases in Alzheimer’s disease is worthy of significant further investigation.

**Figure 4.**
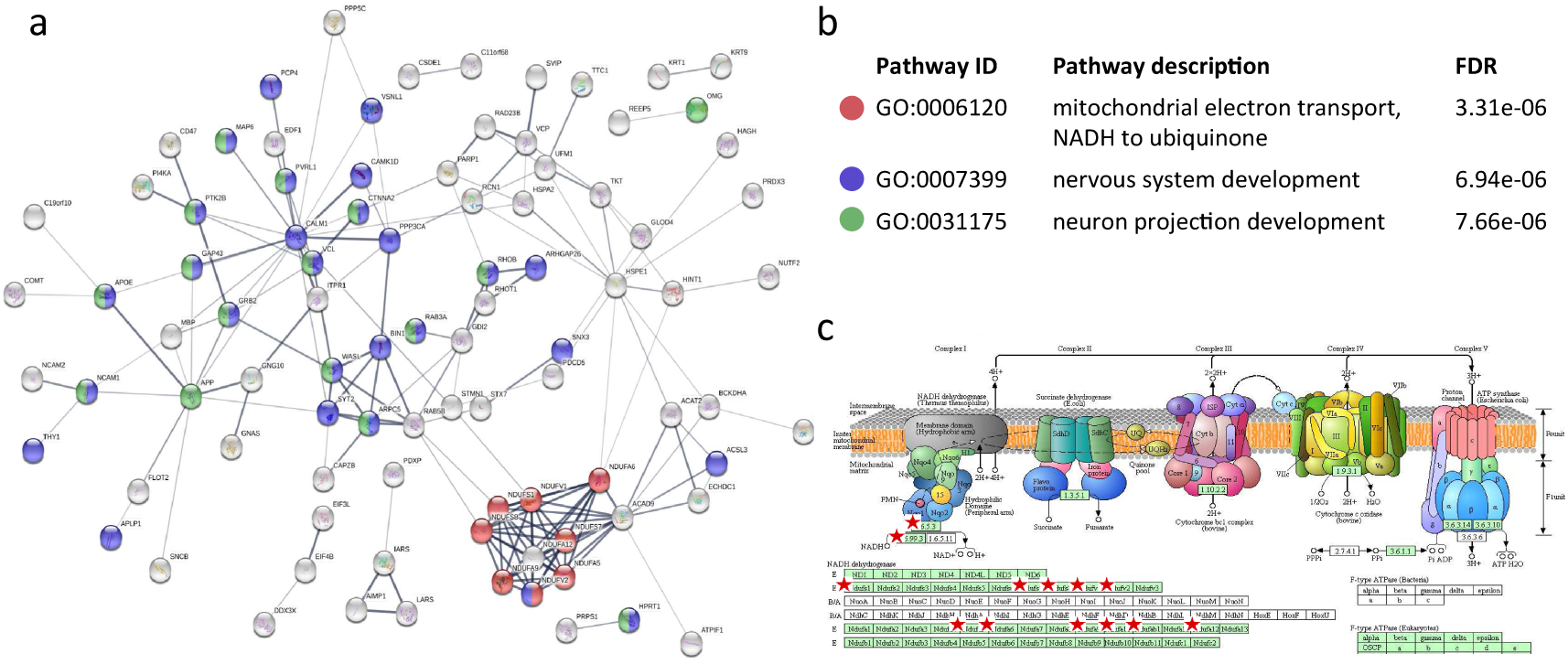
CB-specific biological processes in AD brain. a) 120 proteins that showed CB-specific alterations were enriched for molecular processes in STRING using default setting. Each node represents a protein, and proteins involved in b) significantly enriched pathways were highlighted. c) Dysregulation of the mitochondrial electron transport chain was highlighted by pathway analysis, and proteins affected mapped (red star) into the NADH dehydrogenase complex in KEGG oxidative phosphorylation map.

One of the most distinct changes observed in this CB-specific analysis was that a much greater number of proteins of electron transport chain (ETC) complex 1 were consistently more reduced in abundance (Fig.4b, c Supplementary Data Table 5) than was found in other areas. Furthermore, CB showed increases in oxidative defense proteins involved in glutathione redox reactions and ascorbate recycling (Fig.3c). These data provide strong additional evidence for a protective mechanism in CB that decreases ROS-production by ETC while simultaneously increasing ROS defenses. Another interesting observation in CB was the activation of a Purine Ribonucleosides Degradation pathway, which could not only contribute substrate to the pentose phosphate pathway, but also participate in guanine/guanosine production in this brain region. Combined with the observed activation of Guanine and Guanosine Salvage I pathway, and an increase in guanosine level in CB as previously reported by our metabolomics analysis^14^, these changes may also confer a previously unknown neuroprotective effect in this brain region^43^.

It is well established that CB does not display extensive apoptotic activation seen elsewhere in the brain in Alzheimer’s disease, which is unsurprising given its structurally unaffected status. Our findings indicate that the lack of significant neurodegeneration in this region is not merely due to the absence of an apoptotic signal (e.g. Tau tangles) but instead that CB actively induces a unique pattern of upregulated neuronal survival pathways alongside protection against oxidative and inflammatory damage; a protective mechanism of gene/protein expression which limits disease-related degeneration in this region.

Given the apparently similarity in protein expression which we seen wining each group (severely affected and less affected), we next attempt to identify key regulators of what appears to be a coordinated alteration in protein expression across the brain in response to AD. We performed a correlation network analysis to identify key nodes which may be responsible for the programme of protein expression observed, using the Cytoscape ModuLand plug-in^44^. The resulting correlation network is shown in Figure 5a. Each cluster is coloured differently according to a distinct meta-node, the key regulators of which can be determined by visualizing higher levels of this hierarchy (Fig. 5b). Using this method, we can identify the most influential genes in this correlation network which we hypothesize to be key regulators of protein expression during the pathogenesis of AD. It is noteworthy that in this correlation matrix we are aiming to correlate what we believe to be two distinct processes – AD pathogenesis (seen in HP, ENT, CG, MCx and SCx) and a protective programme that we observe in CB. By overlaying protein expression data onto this network, we can identify which nodes are associated with which process. This overlay (Fig. 5c-h) clearly demonstrates that the correlation network is mainly constructed from proteins involved in AD pathogenesis in the affected regions – few proteins in the network are changed in CB despite the relatively large number of CB proteins which we observe to be changed in the complete dataset. This is to be expected as CB-specific protein changes have limited correlation to the remainder of the dataset. This network is therefore likely to provide a good representation of the key events in AD pathogenesis, and reveals four proteins with the most overall influence on the correlated expression networks: STXBP1 (syntaxin binding protein 1); CRMP1, (collapsin response-mediator protein 1); ACTR10, (actin-related protein 10 homologue); and AMPH (amphiphysin).

**Figure 5.**
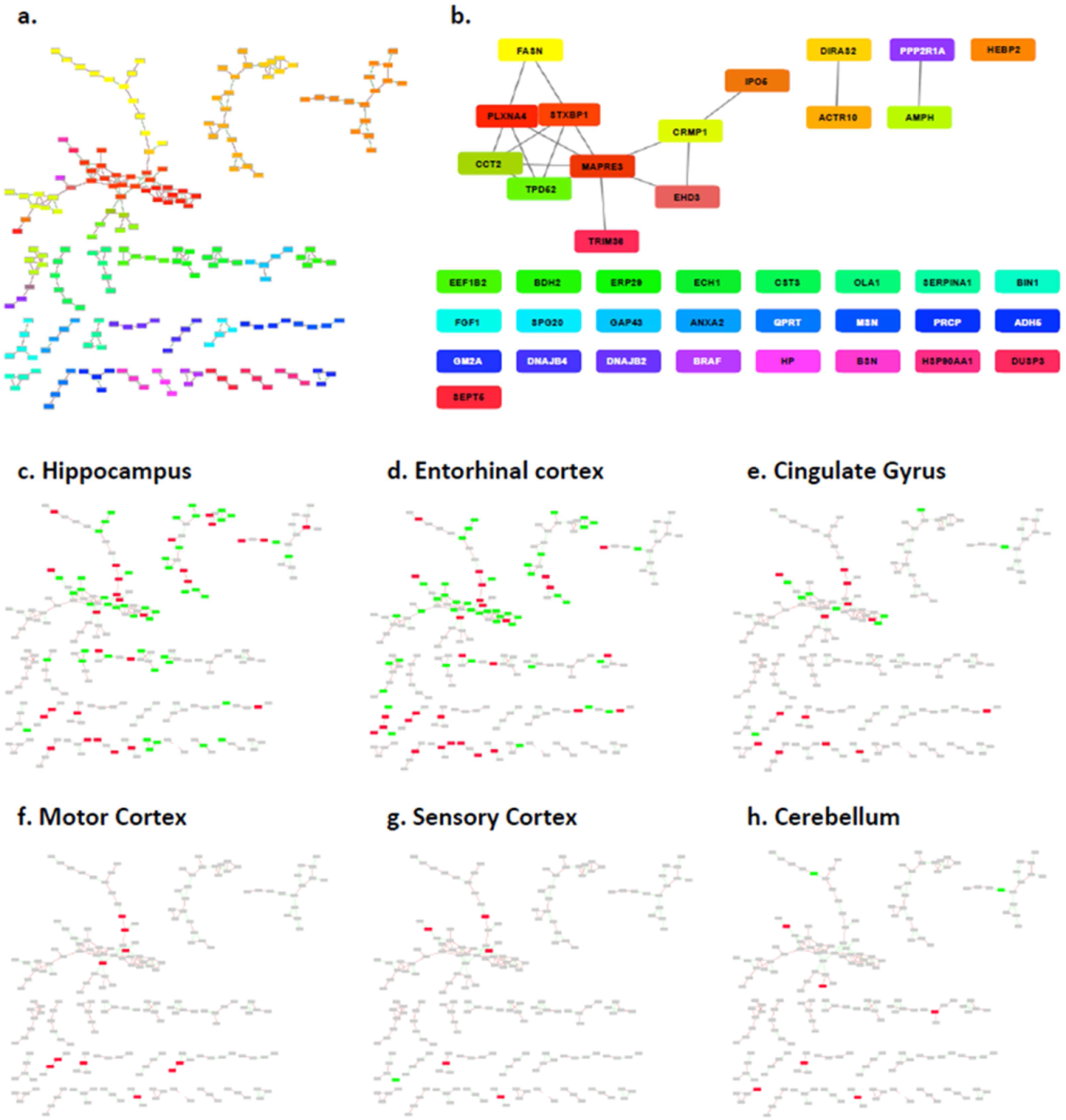
**Global networks analysis** was performed using Cytoscape ModuLand plug-in. a) Correlation network of altered proteins in AD brain, with differently coloured clusters representing different meta-nodes b) key regulators of each meta-node c-h) Overlays of protein expression data from each region and the correlation network.

STXBP1 is the regulator with the most influence in this network. It is reportedly upregulated in AD^45^, has been linked to NFTs^46^ and may interact with PS1^47^. It also plays a major role in neurotransmitter release. STXBP1 thus provides a potential mechanistic explanation for our observation that pathways of neurotransmitter metabolism including dopamine-, noradrenaline-, and serotonin-related signalling showed significant changes in severely affected regions and SCx, but not in MCx or CB. Another important regulator of the network, CRMP1, is part of the semaphorin signalling pathway which is known to guide axons in developing nervous tissue and participates in shaping of neural circuits^48^. ACTR10 may affect prion susceptibility through its involvement in prion propagation and clearance^49^, and has been identified by large scale computational network analyses as one of a large number of potentially important genes in hippocampal ageing, but our finding is novel in AD^50^. The 4^th^ key network regulator identified here, AMPH, is a candidate AD risk gene that may participate in receptor-mediated endocytosis and hence be involved in APP metabolism/clearance^51^. Our finding that these four genes appear to be central to various pathological processes known to be involved in AD development is important, and suggests that further work should be performed to focus on the role of these potentially key mediators of Alzheimer’s disease progession.

Since one of the key factors in AD pathogenesis is thought to be the build-up of amyloid consisting of Aβ peptide generated as a proteolytic product of the Amyloid precursor protein (APP) we examined our data for information about the levels and distribution of these molecules. We found no marked change in APP levels overall but significantly elevated Aβ peptide levels (Supplementary Figure 2a-b), consistent with previous reports^52^. The extent of the increase in Aβ between regions does not appear to follow a gradient of ‘affectedness’, albeit there may be a more pronounced increase in hippocampus. There is no way to determine the primary structure of the Aβ peptide(s) present in each region from these data. Interestingly, while in the AD group almost all samples showed uniformly high levels of Aβ peptide, there was marked variation in levels in control samples (Supplementary Figure 2c). While the quantification of Aβ is necessarily from one peptide, these data emanate from between 5 and 12 unique spectra in each sample we consider this observation is likely robust. This variability is therefore likely to be due to inherent variations in the control population. Although all patients in this group were asymptomatic, it is likely that varying degrees of prodromal disease could have been present, given their age. This is most noticeable in our control 115. While initially assigned as a control, a pathological re-examination performed as a result of the findings of this study and our previous metabolomics analyses^14^ re-classified this individual as a Braak II pre-clinical AD patient. This patient has the highest level of Aβ of all of the control samples and interestingly appears to demonstrate some AD-related changes both in their metabolome and in some of the proteins which we observe to be changed in symptomatic disease. This observation supports the idea that increases in Aβ levels may reflect varying degrees of prodromal disease in these elderly controls. It also demonstrates that studies of the type performed here in earlier stage presymptomatic patients will be critical to further tease out the very earliest events in AD pathogenesis.

In summary, this study provides a map of molecular changes that are present in human *post-mortem* brain tissue in patients with AD and matched controls, providing insights into the brain region specificity of disease at two levels; individual proteins and pathways. We observed global perturbation of protein expression in all six regions of the AD brain which we studied. An association between extent of molecular changes and affectedness was observed for five regions, allowing us to delineate probably ‘early’ and ‘late’ changes in protein expression and revealing previously novel involvement of several pathways and processes. The sixth region, CB, showed an unexpectedly distinct pattern of protein changes, suggestive of induction of a protective response. Correlation network analysis identified four candidate genes STXBP1, CRMP1, ACTR10, and AMPH which may underpin significant portions of the protein expression response to AD. Finally, we recognize that these data have significant value to the community and that other researchers will no doubt wish to assess the status of other AD-related changes not discussed here. As such we have provided all results in an accessible format via a freely-available, searchable on-line database, to allow others to probe specific pathways or individual proteins and their expression in regions across the human Alzheimer’s disease brain and matched controls.

**Supplementary Information** is available in the online version of this paper

## Acknowledgements

The authors would like to thank the families and patients who supported this research by donation of brains to the New Zealand Neurological Foundation Human Brain Bank. This work was supported by Alzheimer’s Research UK (ARUK-PPG2014B-7), the New Zealand Neurological Foundation, the Maurice Wilkins Centre for Molecular Biodiscovery (Tertiary Education Commission 9431-48507; and Doctoral Scholarship funding to Jingshu Xu), the University of Auckland (Doctoral Student funding to Jingshu Xu - JXU058 PReSS) and was facilitated by the Manchester Biomedical Research Centre and the Greater Manchester Comprehensive Local Research Network.

## Author Contributions

J.X., S.P., N.R., I.R-G., B.D.H, and H.W. performed the experiments presented in this manuscript. Initial Bayesian data analysis was developed by A.M.P., A.W.D. and R.D.U. R.H. and P.B. build the web-based data resource. J.X., A.S. and R.D.U. performed data interpretation and network analysis. R.L.M.F., G.J.S.C. and R.D.U. supervised the project. All authors wrote the manuscript.

## Author information

The authors declare no competing financial interests. Correspondence and request for materials, methods or data should be addressed to R.U.

## References

1. Braak, H. & Braak, E. Evolution of neuronal changes in the course of Alzheimer’s disease. Journal of neural transmission. Supplementum 53, 127–140 (1998).

2. Dickson, D. W. Neuropathological diagnosis of Alzheimer’s disease: a perspective from longitudinal clinicopathological studies. Neurobiol Aging 18, S21–26 (1997).

3. Mattson, M. P. Pathways towards and away from Alzheimer’s disease. Nature 430, 631–639, doi:10.1038/nature02621 (2004).

4. Ferreira, I. L., Resende, R., Ferreiro, E., Rego, A. C. & Pereira, C. F. Multiple defects in energy metabolism in Alzheimer’s disease. Current drug targets 11, 1193–1206 (2010).

5. Rudy, C. C., Hunsberger, H. C., Weitzner, D. S. & Reed, M. N. The role of the tripartite glutamatergic synapse in the pathophysiology of Alzheimer’s disease. Aging and disease 6, 131–148, doi:10.14336/ad.2014.0423 (2015).

6. Chen, K. H., Reese, E. A., Kim, H. W., Rapoport, S. I. & Rao, J. S. Disturbed neurotransmitter transporter expression in Alzheimer’s disease brain. Journal of Alzheimer’s disease: JAD 26, 755–766, doi:10.3233/jad-2011-110002 (2011).

7. Webster, S. J., Bachstetter, A. D., Nelson, P. T., Schmitt, F. A. & Van Eldik, L. J. Using mice to model Alzheimer’s dementia: an overview of the clinical disease and the preclinical behavioral changes in 10 mouse models. Frontiers in Genetics 5, 88, doi:10.3389/fgene.2014.00088 (2014).

8. Lu, R. et al. Systems-level dynamic analyses of fate change in murine embryonic stem cells. Nature 462, 358–362, doi:10.1038/nature08575 [doi] (2009).

9. Hawkridge, A. M. & Muddiman, D. C. Mass spectrometry-based biomarker discovery: toward a global proteome index of individuality. Annual review of analytical chemistry (Palo Alto, Calif.) 2, 265–277, doi:10.1146/annurev.anchem.1.031207.112942 (2009).

10. Andreev, V. P. et al. Label-free quantitative LC-MS proteomics of Alzheimer’s disease and normally aged human brains. J Proteome Res 11, 3053–3067, doi:10.1021/pr3001546 (2012).

11. Hondius, D. C. et al. Profiling the human hippocampal proteome at all pathologic stages of Alzheimer’s disease. Alzheimer’s & Dementia 12, 654–668, doi:http://dx.doi.org/10.1016/j.jalz.2015.11.002 (2016).

12. Musunuri, S. et al. Quantification of the brain proteome in Alzheimer’s disease using multiplexed mass spectrometry. J Proteome Res 13, 2056–2068, doi:10.1021/pr401202d (2014).

13. Manavalan, A. et al. Brain site-specific proteome changes in aging-related dementia. Experimental & molecular medicine 45, e39, doi:10.1038/emm.2013.76 (2013).

14. Xu, J. et al. Graded perturbations of metabolism in multiple regions of human brain in Alzheimer’s disease: Snapshot of a pervasive metabolic disorder. Biochimica et biophysica acta 1862, 1084–1092, doi:10.1016/j.bbadis.2016.03.001 (2016).

15. Braak, H., Braak, E., Bohl, J. & Lang, W. Alzheimer’s disease: amyloid plaques in the cerebellum. J Neurol Sci 93, 277–287 (1989).

16. Wegiel, J. et al. Cerebellar atrophy in Alzheimer’s disease-clinicopathological correlations. Brain Res 818, 41–50 (1999).

17. Dukart, J. et al. Differential effects of global and cerebellar normalization on detection and differentiation of dementia in FDG-PET studies. NeuroImage 49, 1490–1495, doi:10.1016/j.neuroimage.2009.09.017 (2010).

18. Lyoo, C. H. et al. Cerebellum Can Serve As a Pseudo-Reference Region in Alzheimer Disease to Detect Neuroinflammation Measured with PET Radioligand Binding to Translocator Protein. Journal of Nuclear Medicine 56, 701–706, doi:10.2967/jnumed.114.146027 (2015).

19. Guo, C. C. et al. Network-selective vulnerability of the human cerebellum to Alzheimer’s disease and frontotemporal dementia. Brain 139, 1527–1538, doi:10.1093/brain/aww003 (2016).

20. Ishii, K. et al. Reduction of cerebellar glucose metabolism in advanced Alzheimer’s disease. Journal of nuclear medicine: official publication, Society of Nuclear Medicine 38, 925–928 (1997).

21. Xu, J. et al. Elevation of brain glucose and polyol-pathway intermediates with accompanying brain-copper deficiency in patients with Alzheimer’s disease: metabolic basis for dementia. Scientific reports 6, 27524, doi:10.1038/srep27524 (2016).

22. Ivakhno, S. & Armstrong, J. D. Non-linear dimensionality reduction of signaling networks. BMC systems biology 1, 27, doi:10.1186/1752-0509-1-27 (2007).

23. Ray, M. & Zhang, W. Analysis of Alzheimer’s disease severity across brain regions by topological analysis of gene co-expression networks. BMC systems biology 4, 136, doi:10.1186/1752-0509-4-136 (2010).

24. Heneka, M. T., Golenbock, D. T. & Latz, E. Innate immunity in Alzheimer’s disease. Nature immunology 16, 229–236, doi:10.1038/ni.3102 (2015).

25. Lovestone, S. et al. AddNeuroMed--the European collaboration for the discovery of novel biomarkers for Alzheimer’s disease. Ann N Y Acad Sci 1180, 36–46, doi:10.1111/j.1749-6632.2009.05064.x (2009).

26. Harold, D. et al. Genome-wide association study identifies variants at CLU and PICALM associated with Alzheimer’s disease. Nature genetics 41, 1088–1093, doi:10.1038/ng.440 (2009).

27. Lambert, J. C. et al. Genome-wide association study identifies variants at CLU and CR1 associated with Alzheimer’s disease. Nature genetics 41, 1094–1099, doi:10.1038/ng.439 (2009).

28. Zhang, D. F. et al. CFH Variants Affect Structural and Functional Brain Changes and Genetic Risk of Alzheimer’s Disease. Neuropsychopharmacology: official publication of the American College of Neuropsychopharmacology 41, 1034–1045, doi:10.1038/npp.2015.232 (2016).

29. Van Eldik, L. J. et al. The roles of inflammation and immune mechanisms in Alzheimer’s disease. Alzheimer’s & Dementia: Translational Research & Clinical Interventions 2, 99–109, doi:https://doi.org/10.1016/j.trci.2016.05.001 (2016).

30. Xu, Y., Stamenkovic, I. & Yu, Q. CD44 attenuates activation of the Hippo signaling pathway and is a prime therapeutic target for glioblastoma. Cancer research 70, 2455–2464, doi:10.1158/0008-5472.CAN-09-2505 (2010).

31. Kapogiannis, D. & Mattson, M. P. Disrupted energy metabolism and neuronal circuit dysfunction in cognitive impairment and Alzheimer’s disease. The Lancet. Neurology 10, 187–198, doi:10.1016/s1474-4422(10)70277-5 (2011).

32. Bradley, K. M. et al. Cerebral perfusion SPET correlated with Braak pathological stage in Alzheimer’s disease. Brain 125, 1772–1781 (2002).

33. Arnaiz, E. et al. Impaired cerebral glucose metabolism and cognitive functioning predict deterioration in mild cognitive impairment. Neuroreport 12, 851–855 (2001).

34. Costantini, L. C., Barr, L. J., Vogel, J. L. & Henderson, S. T. Hypometabolism as a therapeutic target in Alzheimer’s disease. BMC neuroscience 9 Suppl 2, S16, doi:10.1186/1471-2202-9-s2-s16 (2008).

35. Yao, J., Rettberg, J. R., Klosinski, L. P., Cadenas, E. & Brinton, R. D. Shift in brain metabolism in late onset Alzheimer’s disease: implications for biomarkers and therapeutic interventions. Molecular aspects of medicine 32, 247–257, doi:10.1016/j.mam.2011.10.005 (2011).

36. Foxton, R. H., Land, J. M. & Heales, S. J. Tetrahydrobiopterin availability in Parkinson’s and Alzheimer’s disease; potential pathogenic mechanisms. Neurochemical research 32, 751–756, doi:10.1007/s11064-006-9201-0 (2007).

37. Eixarch, H., Gutiérrez-Franco, A., Montalban, X. & Espejo, C. Semaphorins 3A and 7A: potential immune and neuroregenerative targets in multiple sclerosis. Trends in Molecular Medicine 19, 157–164, doi:10.1016/j.molmed.2013.01.003.

38. Lee, J. W. et al. Editing-defective tRNA synthetase causes protein misfolding and neurodegeneration. Nature 443, 50–55, doi:10.1038/nature05096 (2006).

39. Ishimura, R. et al. Ribosome stalling induced by mutation of a CNS-specific tRNA causes neurodegeneration. Science (New York, N.Y.) 345, 455–459, doi:10.1126/science.1249749 (2014).

40. Tshori, S., Razin, E. & Nechushtan, H. Amino-acyl tRNA synthetases generate dinucleotide polyphosphates as second messengers: functional implications. Topics in current chemistry 344, 189–206, doi:10.1007/128_2013_426 (2014).

41. Ding, Q., Markesbery, W. R., Chen, Q., Li, F. & Keller, J. N. Ribosome Dysfunction Is an Early Event in Alzheimer’s Disease. The Journal of Neuroscience 25, 9171 (2005).

42. Guo, M. & Schimmel, P. Essential nontranslational functions of tRNA synthetases. Nature chemical biology 9, 145–153, doi:10.1038/nchembio.1158 (2013).

43. Lanznaster, D., Dal-Cim, T., Piermartiri, T. C. B. & Tasca, C. I. Guanosine: a Neuromodulator with Therapeutic Potential in Brain Disorders. Aging and disease 7, 657–679, doi:10.14336/AD.2016.0208 (2016).

44. Szalay-Beko, M. et al. ModuLand plug-in for Cytoscape: determination of hierarchical layers of overlapping network modules and community centrality. Bioinformatics (Oxford, England) 28, 2202–2204, doi:10.1093/bioinformatics/bts352 (2012).

45. Donovan, L. E. et al. Analysis of a membrane-enriched proteome from postmortem human brain tissue in Alzheimer’s disease. Proteomics Clin Appl 6, 201–211, doi:10.1002/prca.201100068 (2012).

46. Takahashi, M., Iseki, E. & Kosaka, K. Cdk5 and munc-18/p67 co-localization in early stage neurofibrillary tangles-bearing neurons in Alzheimer type dementia brains. J Neurol Sci 172, 63–69 (2000).

47. Inoue, M. et al. Human brain proteins showing neuron-specific interactions with gammasecretase. The FEBS journal 282, 2587–2599, doi:10.1111/febs.13303 (2015).

48. Uesaka, N. et al. Retrograde semaphorin signaling regulates synapse elimination in the developing mouse brain. Science 344, 1020 (2014).

49. Brown, C. A. et al. In vitro screen of prion disease susceptibility genes using the scrapie cell assay. Human molecular genetics 23, 5102–5108, doi:10.1093/hmg/ddu233 (2014).

50. Dempsey, K. M. & Ali, H. H. Identifying aging-related genes in mouse hippocampus using gateway nodes. BMC systems biology 8, 62, doi:10.1186/1752-0509-8-62 (2014).

51. Talwar, P. et al. Genomic convergence and network analysis approach to identify candidate genes in Alzheimer’s disease. BMC genomics 15, 199, doi:10.1186/1471-2164-15-199 (2014).

52. Seyfried, N. T. et al. A Multi-network Approach Identifies Protein-Specific Co-expression in Asymptomatic and Symptomatic Alzheimer’s Disease. Cell systems 4, 60-72.e64, doi:10.1016/j.cels.2016.11.006 (2017).

