## Supplementary Materials for "Regional protein expression in human Alzheimer’s brain correlates with disease severity"

### Methods

***Human brains.*** All experiments were performed in accordance with relevant guidelines and regulations. The case-control study of *post-mortem* human brain was approved by the University of Auckland Human Participants Ethics Committee with informed consent from all families.

Human brains were obtained from the New Zealand Neurological Foundation Human Brain Bank, University of Auckland [1]. Each brain was dissected under the supervision of neuroanatomists (JX, SP, HJW and RLMF), who accurately identified each region. Brain regions studied were hippocampus, entorhinal cortex, cingulate gyrus, sensory cortex, motor cortex and cerebellum: grey matter from each region was sampled. Aliquots of 100±5 mg were dissected from each region and stored at -80°C until analysis, and were otherwise treated as previously described [2]. Patients had *ante-mortem* evidence of clinical dementia, whereas controls did not. Controls were selected by matching for age, sex and *post-mortem* delay. A consultant neuropathologist diagnosed or excluded AD by applying the Consortium to Establish a Registry for Alzheimer's Disease (CERAD) criteria [3], and determined the neuropathological severity by assigning the Braak stage [4] and amyloid load by applying the 2013 consensus National Institute on Aging–Alzheimer's Association guidelines [5] (Supplementary Data Table1). One control patient (115) had neuropathological findings consistent with AD (Braak Stage II; Supplementary Data Table 1) and was therefore diagnosed with preclinical disease: this finding is consistent with the known frequency of asymptomatic AD in similarly-aged groups in the study population [6].

***Protein extraction and preparation for iTRAQ.*** Protein extraction and preparation for iTRAQ was carried out according to a previously described method [7], with each brain region analysed independently. Brain tissue samples of 100±5 mg were extracted in 500 µL 1 M Triethylammonium bicarbonate buffer (TEAB) + 0.1% (w/v) SDS, and homogenized at 25 Hz (2 × 3 min) with a Qiagen tissuelyser. The tubes were then vortexed for 10 s and centrifuged at 4°C for 5 min at 13,400 × g. The supernatants were transferred into a new set of tubes and protein concentration was determined by using Bradford protein assay (Bio-Rad Protein Assay Dye Reagent Concentrate) and a SpectraMax M5 plate-reader (Molecular Devices). From each sample, a volume equivalent to 100 µg protein was transferred into a new set of tubes for further processing. Identical reference pool samples (total of 100 µg protein per reference sample) were made by combining portions from four representative individual samples from each group, AD and control. All samples were equalised for final volume using 1 M TEAB + 0.1% (w/v) SDS.

Protein samples were reduced by addition of 0.1 volume of 50 mM DTT, followed by incubation at 60 °C for 30 min. Alkylation was carried out by addition of 0.05 volumes of 200 mM iodoacetamide, followed by incubation in the dark at room temperature for 10-15 min. Protein digestion was subsequently carried out overnight at 37°C, by adding 10 μg of modified porcine trypsin (Promega) resuspended in 1 M TEAB, ensuring the final SDS concentration fell below 0.05% (w/v). After digestion, the samples were dried completely in an Eppendorf concentrator, and re-suspended in 30 μL 1 M TEAB to achieve equal volume across all samples before iTRAQ labelling.

The iTRAQ labelling was carried out according to the manufacturer's instruction using the 8-plex iTRAQ kit (AB Sciex). Briefly, vials containing iTRAQ reagent were thawed on the bench for 2-3 min. After spinning the samples down, 70 μL isopropanol was added to each vial, followed by a pulse spin. The content of the vials was then transferred to the protein samples and then incubated on the bench for 2-3 h. Each 8-plex contained two separate digests of the reference pool sample, three AD samples and three control samples.

iTRAQ-labelled samples destined for the same LC-MS/MS run were pooled, followed by a spin at 13,400 × g for 5 min. Each pooled sample was then divided into 2 equal aliquots and dried completely using an Eppendorf centrifugal evaporator concentrator. One pooled aliquot from each 8-plex experiment was subjected to high-pH reverse-phase (HpHRP) for peptide fractionation. Remaining dried-pool aliquots were stored at -80 $^{o}$C for repeated analysis if required.

***High-pH reversed-phase fractionation (HpHRP).*** HpHRP was performed using an Agilent HPLC 1200 system (Agilent, Santa Clara, California). Reversed-phase chromatography buffers (buffer A; 0.1% (v/v) ammonium hydroxide in HPLC-grade water and buffer B; 0.1% (v/v) ammonium hydroxide in acetonitrile) were made fresh.

Each iTRAQ-labelled pool sample was resuspended in 900 μL of 3% (v/v) buffer B and loaded onto a high-pH reversed-phase column (ZORBAX 300Extend-C18 4.6 × 150mm 3.5micron, Agilent) for 40-min with a flow of 1mL/min at 3% (v/v) buffer B. The peptides were then eluted using the gradient as follows (minutes:%B); 0:3, 5:3, 30:27, 35:50, 36:100, 41:100, 42:3. A total of 86 fractions were collected in a 96-well plate, which was dried in a centrifugal evaporator (Eppendorf) and stored at –20 °C prior to LC-MS/MS analysis.

***Low-pH LC-Mass spectrometry based analysis.*** Each fraction was re-suspended in 27 μL of 97% water + 3% acetonitrile + 0.1% trifluoroacetic acid (TFA; v/v/v) and 9 μL was injected into a nano-Acquity UPLC system (Waters). Peptides were trapped on a nanoACQUITY 2G-V/M Trap Sym C18 5 μm 180 μm × 20 mm (Waters) and washed at a flow rate of 7.5 μL/min for 10 min. Peptides were then eluted and chromatographed using a nanoACQUITY BEH300

C18 1.7 µm 75 µm × 250 mm (Waters) at 300 nL/min using following gradient profile (minutes:%B); 0:3, 3:3, 91:40, 93:90, 108:90, 109:3, 130:3. The buffers used were: buffer A: 97% water + 3% acetonitrile + 0.1% formic acid and buffer B: 100% acetonitrile + 0.1% formic acid (v/v).

The eluent was directed into an ESI microionspray II source of a QSTAR Elite Q-TOF spectrometer (AB SCIEX) scanning in MS from 400 to 1200 m/z. Multiply-charged peptides (2+ to 4+) were selected for MS/MS analysis (110–1200 m/z). The information-dependent acquisition (IDA) settings were: 4 precursors per cycle and cycle times (MS 0.75 s, MS/MS1 0.75 s, MS/MS2 0.75 s, MS/MS3 1 s and MS/MS4 1 s). Selected peptides were fragmented twice and then dynamically excluded for 90 s. The resulting data were searched against the human component of the Swissprot database (release 2013_03) using Protein-Pilot v4.0 (AB SCIEX). Search parameters were: iTRAQ 8plex, trypsin; cys alkylation, iodoacetamide; search effort, thorough. A total of 40466 proteins were searched. To perform false discovery rate analysis on the protein identification, the search database was reversed and concatenated with the forward database and used as the search DB within ProteinPilot. False discovery rate was determined by calculating the number of reverse 'hits' as a proportion of 'forward' hits using the dedicated worksheet exported from the search software.

***Data processing.*** Bayesian protein-level differential quantification was performed separately for each brain region using v1.0.0 of the in-house developed software 'BayesProt' (https://github.com/biospi/bayesprot/releases/tag/v1.0.0). An earlier version of this technique was presented in Freeman et al. [8], which combined Protein-Pilot (AB SCIEX) sample normalization ('bias correction') with a Bayesian linear mixed-effects model implemented with the MCMCglmm R Package [9]. Analysis of each brain region in isolation adds strength to our comparison of protein expression changes across multiple regions, as these were identified and quantified independently.

Since iTRAQ measurements from Time-of-Flight instruments are recorded as discrete ion counts, and technical/biological variation are assumed log-normal, we adopted a Generalised Linear Mixed Model (GLMM) with Poisson likelihood and log-link, where each protein was modelled separately using peptide measurements unique to that protein. The sample normalization factors represent the mass spectrometer's exposure to each sample, and hence were included as a fixed offset within the model. The current version of BayesProt additionally (i) enables estimation of both biological and digestion variance through the incorporation of multiple digests for a single sample (i.e. the six reference pool digests), (ii) negates the need for Protein-Pilot normalization by implementing a two-stage GLMM and (iii) provides a simplified Markov Chain Monte Carlo (MCMC) mixing criterion for both stages.

In both stages: (a) for each peptide a separate random digest effect is fitted, which has the effect of weighting each peptide's contribution to the protein-level quantification by its reproducibility across digests; (b) the set of measurement channels within each iTRAQ spectrum are each assigned (i) a baseline fixed effect to account for varying selection/ionisation/fragmentation efficiencies across spectra, and (ii) an independent log-normal residual variance to account for over-dispersion due to background contamination and incorrectly identified spectra. In stage one, we also model the interaction between LC-MS/MS run and iTRAQ channel as a fixed effect *i.e.* within each run, we infer the protein-level log ratio between iTRAQ channel 113 and channels 114, 115, 116, 117, 118, 119 & 121. For each channel relative to 113, the result is a set of posterior probability distributions, one for each protein in the study; these are combined to derive a posterior distribution for the median log ratio for each channel relative to 113, which is taken as the inferred sample normalization factors.

In stage two, rather than using point estimates of the normalization factors as fixed sample offsets, a set of sample fixed effects are fitted which have prior distributions set to the means and variances of the inferred median log ratio distributions. In addition, in stage two we specify the full experimental design: (a) protein-level differential expression fold change between cases and controls is fitted as a condition fixed effect (with control as baseline); (b) to unequal biological variance across cases and controls, subject is treated as two random effects, one for control samples and one for cases. Using the inferred posterior distribution of the condition fixed effect, we performed a one-sided significance test on the posterior probability that the mean fold-change is either above or below ±1.05 – i.e. at least a 5% change from control – denoted as P (1.05fc). The reciprocal of this posterior probability represents the local False Discovery Rate (lFDR), the probability that this specific test is a false discovery. In this study, we defined significance using a *global* FDR threshold of 5% i.e. the largest set of proteins with an average lFDR ≤5% were deemed significant and hence delivered to downstream pathway analysis. The condition fixed effect posterior distributions, FDRs and descriptive statistics (mean log ratio plus 95% highest posterior density interval) for every protein across all regions are presented online (www.manchester.ac.uk/dementia-proteomes-project). Posterior distributions of per-sample protein quantifications are also presented, derived from the latent variables of the sample random effects.

Residual variances were assigned inverse-Gamma priors, while random effects were assigned parameter-expanded Cauchy priors. The model was tested with different prior scale factors to establish that the priors were not informative to the outcome. In stages one and two, the model was run with 10 and 100 MCMC chains per protein, respectively, each chain consisting of 10,000 samples preceded by 3,000 burn-in samples. Mixing was assessed using

Warnes & Raftery's MCGibbsit run-length diagnostic, combining the estimate error-bounding approach of Raftery and Lewis with the between chain variance verses within chain variance approach of Gelman and Rubin (https://cran.r-project.org/web/packages/mcgibbsit/index.html).

For a protein to be considered quantified sufficiently well to be included in downstream pathway, correlation and comparative analyses, we require identification and quantification from at least three spectra. This quality control is important when making comparisons across datasets as it ensures that only high-quality protein quantitation is taken forward into comparative studies, reducing 'noise'.

***Data Analysis*** Processed protein-level data were analysed through a range of software tools. Data alignment, filtering and characterisation was initially performed in Microsoft Excel. Heat maps were constructed using Cluster 3.0 (http://bonsai.hgc.jp/~mdehoon/software/cluster/software.htm) and viewed using Java TreeView (https://sourceforge.net/projects/jtreeview/files/; [10]). Venn diagrams were built using the Interactive Venn tool (www.interactivenn.net; [11]). The Isomap algorithm [12, 13] was implemented in Qlucore Omics Explorer (version 3.2, Qlucore, Lund, Sweden).

***Network Analysis.*** Pathway enrichment analysis was performed for each brain region independently using Ingenuity Pathway Analysis (Qiagen), selecting the user dataset as the background and considering only relationships which had been experimentally observed or predicted with 'high' confidence, and was limited to human interactions. Following analysis, significant pathways were reviewed and those which contained genes which formed a complete subset of another pathway were removed. In parallel, we performed functional annotation clustering analysis for each brain region using online DAVID (https://david.ncifcrf.gov/; [14, 15]) with custom classification stringency setting; similarity term overlap=5, similarity threshold=0.95, initial group membership=3, final group membership=3, multiple linkage threshold=0.5, EASE=1.0, and Benjamini correction. Enrichment score ≥1.3 was considered significant and highlighted in supplementary table 5. Protein-protein interaction networks were analysed for proteins that were uniquely altered in CB only, using online STRING (https://string-db.org; [16]) at default setting.

To identify key regulators of protein expression, a correlation matrix of protein expression across the six brain tissue samples was generated in Qlucore Omics Explorer and modified in R to only contain proteins with a $|0.9|$ r-value. The network was visualised in Cytoscape

[17]. Protein modules with correlated expression were identified using the Moduland algorithm [18] and arranged in a hierarchy based on their network centrality.

***Data Availability.*** Raw mass spectral data, along with extracted .mgf peaklists and ProteinPilot .group results files are available via the PRIDE data repository, with each brain region submitted independently to reflect the way in which the study was performed. PRIDE accessions are:

| Region | PRIDE AC |
|---|---|
| Hippocampus | PXD008739 |
| Entorhinal cortex | PXD008806 |
| Cingulate gyrus | PXD008779 |
| Motor cortex | PXD008807 |
| Sensory cortex | PXD008753 |
| Cerebellum | PXD008755 |

We recognize that these data require specialist interpretation. To support data sharing we have also made available the outputs of our initial MS analysis (after ProteinPilot database searching for peptide/protein identification and peptide relative quantification) by depositing the Protein Summary (protein identification data) and Peptide Summary (peptide identification data), along with the raw MS peak lists, with the Open Science Framework (https://osf.io), which can be accessed by the following DOI 10.17605/OSF.IO/6BXJQ.

Reference:


1. Waldvogel, H.J., et al., *The collection and processing of human brain tissue for research.* Cell Tissue Bank, 2008. **9**: p. 169-179.
2. Schönberger, S.J., et al., *Proteomic analysis of the brain in Alzheimer's disease: molecular phenotype of a complex disease process.* Proteomics, 2001. **1**(12): p. 1519-1528.
3. Mirra, S.S., et al., *The Consortium to Establish a Registry for Alzheimer's Disease (CERAD). Part II. Standardization of the neuropathologic assessment of Alzheimer's disease.* Neurol, 1991. **41**: p. 479-486.
4. Braak, H. and E. Braak, *Neuropathological stageing of Alzheimer-related changes.* Acta Neuropathol, 1991. **82**: p. 239-259.

5. Montine, T.J., et al., *National Institute on Aging-Alzheimer's Association guidelines for the neuropathologic assessment of Alzheimer's disease: a practical approach.* Acta Neuropathol, 2012. **123**: p. 1-11.
6. Skoog, I., *Detection of preclinical Alzheimer's disease.* N Engl J Med, 2000. **343**(7): p. 502-503.
7. Unwin, R.D., J.R. Griffiths, and A.D. Whetton, *Simultaneous analysis of relative protein expression levels across multiple samples using iTRAQ isobaric tags with 2D nano LC-MS/MS.* Nat Protoc, 2010. **5**(9): p. 1574-82.
8. Freeman, O.J., et al., *Metabolic dysfunction is restricted to the sciatic nerve in experimental diabetic neuropathy.* Diabetes, 2015.
9. Hadfield, J., *MCMC Methods for Multi-Response Generalized Linear Mixed Models: The MCMCglmm R Package.* Journal of Statistical Software, 2010. **33**(2): p. 1-22.
10. Saldanha, A.J., *Java Treeview--extensible visualization of microarray data.* Bioinformatics, 2004. **20**(17): p. 3246-8.
11. Heberle, H., et al., *InteractiVenn: a web-based tool for the analysis of sets through Venn diagrams.* BMC Bioinformatics, 2015. **16**: p. 169.
12. Tenenbaum, J.B., V. de Silva, and J.C. Langford, *A global geometric framework for nonlinear dimensionality reduction.* Science, 2000. **290**(5500): p. 2319-23.
13. Nilsson, J., et al., *Approximate geodesic distances reveal biologically relevant structures in microarray data.* Bioinformatics, 2004. **20**(6): p. 874-80.
14. Huang da, W., B.T. Sherman, and R.A. Lempicki, *Bioinformatics enrichment tools: paths toward the comprehensive functional analysis of large gene lists.* Nucleic Acids Res, 2009. **37**(1): p. 1-13.
15. Huang da, W., B.T. Sherman, and R.A. Lempicki, *Systematic and integrative analysis of large gene lists using DAVID bioinformatics resources.* Nat Protoc, 2009. **4**(1): p. 44-57.
16. Szklarczyk, D., et al., *STRING v10: protein-protein interaction networks, integrated over the tree of life.* Nucleic Acids Res, 2015. **43**(Database issue): p. D447-52.
17. Smoot, M.E., et al., *Cytoscape 2.8: new features for data integration and network visualization.* Bioinformatics, 2011. **27**(3): p. 431-2.
18. Szalay-Beko, M., et al., *ModuLand plug-in for Cytoscape: determination of hierarchical layers of overlapping network modules and community centrality.* Bioinformatics, 2012. **28**(16): p. 2202-4.
