## Supplementary Materials for "Regional protein expression in human Alzheimer’s brain correlates with disease severity"

### Supplementary Information : User guide for navigating online datasets.

To aid in the dissemination and utility of these data, we have established an online repository for this study from which it is possible to search our datasets for specific genes/proteins of interest and determine both their relative quantity, and the quality of that data, in our study. This repository is housed at www.manchester.ac.uk/dementia-proteomes-project.

Here we provide a brief overview of this resource, and how it can be used and how data are interpreted.

Data is searched via a free text search which will search either the gene name, or any word in the protein description, SWISS-Prot ID or accession number.

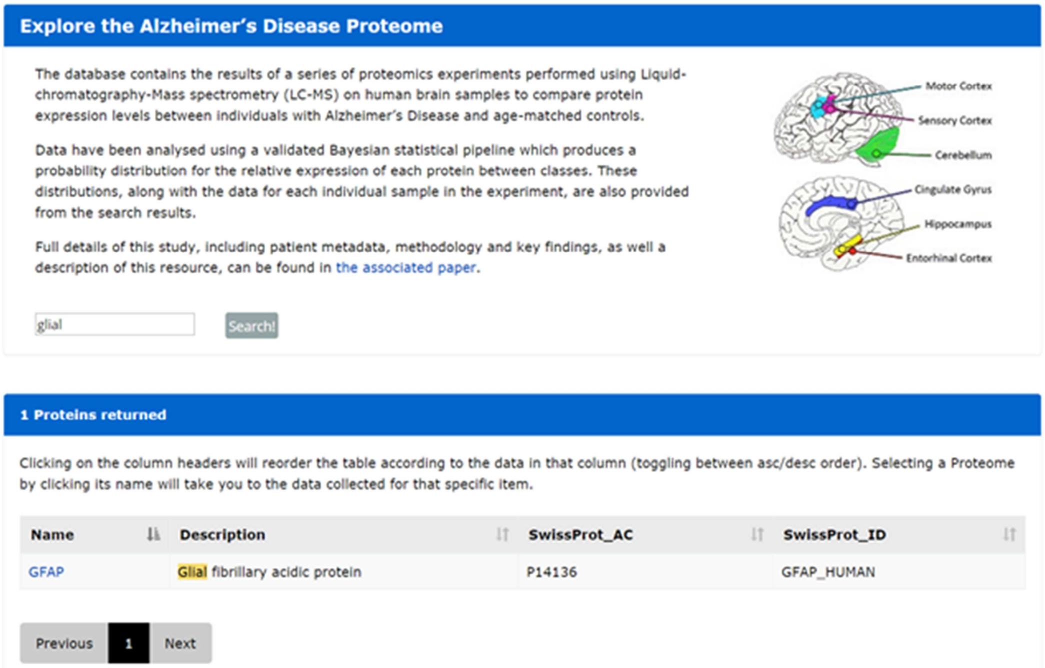

**Explore the Alzheimer's Disease Proteome**

The database contains the results of a series of proteomics experiments performed using Liquid-chromatography-Mass spectrometry (LC-MS) on human brain samples to compare protein expression levels between individuals with Alzheimer's Disease and age-matched controls.

Data have been analysed using a validated Bayesian statistical pipeline which produces a probability distribution for the relative expression of each protein between classes. These distributions, along with the data for each individual sample in the experiment, are also provided from the search results.

Full details of this study, including patient metadata, methodology and key findings, as well a description of this resource, can be found in the associated paper.



glial Search!

**1 Proteins returned**

Clicking on the column headers will reorder the table according to the data in that column (toggling between asc/desc order). Selecting a Proteome by clicking its name will take you to the data collected for that specific item.

| Name | Description | SwissProt_AC | SwissProt_ID |
|---|---|---|---|
| GFAP | Glial fibrillary acidic protein | P14136 | GFAP_HUMAN |

Previous 1 Next

Selecting the Name of the protein of interest opens a table (below) showing the regions for which we have quantification data for that protein, and associated data as follows:

**GFAP**

AC : **P14136**
ID : **GFAP_HUMAN**
Description : **Glial fibrillary acidic protein**

| Brain Area | Peptides | Spectra | Log$_2$(fc) | Lower | Upper | LocalFDR | GlobalFDR |
|---|---|---|---|---|---|---|---|
| Hippocampus (HIP) | 95 | 1829 | 0.529149485 | -0.066167874 | 1.103791236 | 0.0573 | 0.021314341 |
| Motor Cortex (MC) | 44 | 590 | 0.198755814 | -0.247789602 | 0.661911914 | 0.2655 | 0.140215328 |
| Sensory Cortex (SC) | 57 | 821 | 0.095872844 | -0.330819011 | 0.544217367 | 0.448 | 0.259713819 |
| Cerebellum (CER) | 61 | 794 | 0.211085361 | -0.153213647 | 0.570785583 | 0.1952 | 0.087537702 |
| Entorhinal Cortex (ERC) | 70 | 1045 | 0.49231705 | -0.078521237 | 1.084060479 | 0.0681 | 0.028219935 |
| Middle Temporal Gyrus (MTG) | 49 | 901 | 0.524556034 | 0.096108562 | 0.951143024 | 0.0196 | 0.007880435 |
| Cingulate Gyrus (CG) | 54 | 1211 | 0.46881308 | -0.168811081 | 1.094320968 | 0.0961 | 0.040010067 |

**Peptides** – the number of distinct peptides identified associated with the protein of interest.

**Spectra** – the number of distinct MS/MS spectra associated with the protein of interest.

**Log$_2$(fc)** – The Log to base 2 of the fold change (fc) difference between Alzheimer's Disease and Control tissues. Positive numbers imply a relative increase in AD, negative numbers imply a relative decrease.

**Lower and Upper** – Describe the lower and upper 95% credible intervals for the protein relative quantitation in that region.

**LocalFDR and GlobalFDR** – One minus the probability that this protein exhibits at least a 5% deviation from control levels in AD (local FDR) and a calculated global FDR demonstrating the average of localFDRs across the dataset.

To explore these data in more depth, the user can select the Log2(fc) ratio for each condition and view the probability density plot for that specific protein in that region. Note that the algorithm calculates a localFDR (1-p(protein differs from control by at least 5%) both for upregulation and downregulation. However for each protein only the lowest local FDR i.e. most likely result is listed.

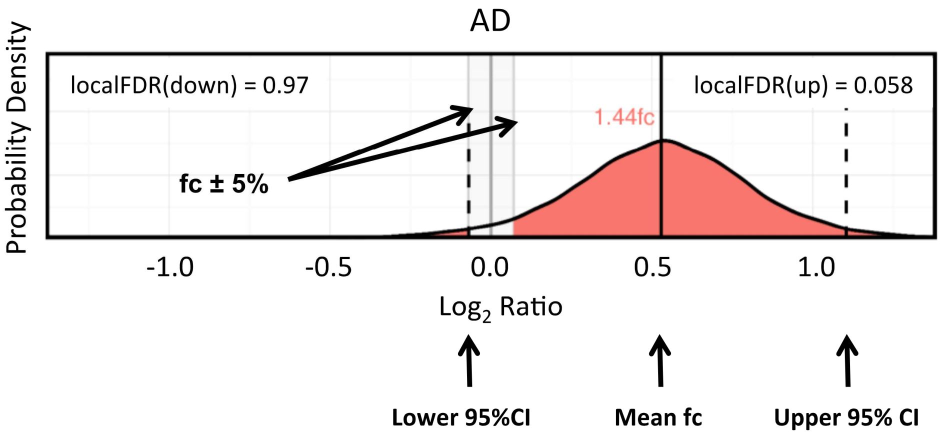


To view variability across all samples in the study, the user can also select the region name. This allows them to view the posterior distribution of each protein on a per-sample basis.

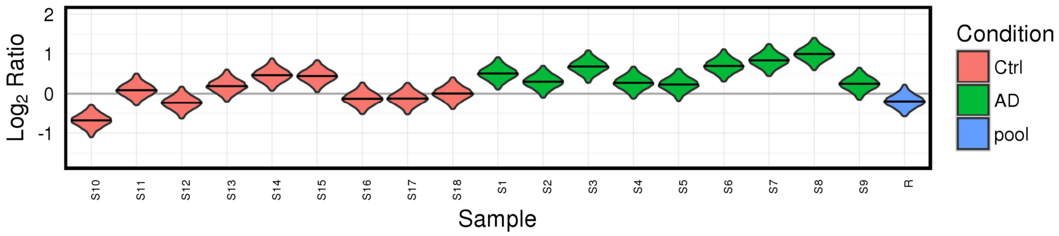
