## Supplementary Materials for "Regional protein expression in human Alzheimer’s brain correlates with disease severity"

### Supplementary Legends

**Supplementary Figure.** Expression probability distributions for a) the Amyloid precursor protein and b) the amyloid beta peptide (sum of 1-40 and 1-42 peptides, which are not distinguishable in this assay) for all brain regions under study. Each plot shows the probability distribution along with the most likely mean expression ratio and the calculated FDR (1-probability) for each molecule having a differential expression between cases and control of at least 5%. c) Bayesian probability distributions for estimated levels of amyloid beta peptides 1-40 plus 1-42 ion each individual sample in this study.

**Supplementary Table 1.** Clinical characteristics of AD and Control brains used for current study. Brain pathology and Braak stage were analysed by a qualified neuropathologist and cause of death was determined at post-mortem examination. Abbreviations: AD, Alzheimer's disease; F, female; M, male. Patient H241 was found to have post-mortem signs consistent with AD and was described as A3, B1, C1 using the 'ABC' criteria for AD neuropathologic change that incorporates histopathological assessments of Aβ deposits (A), staging of neurofibrillary tangles (B), and scoring of neuritic plaques (C).The corresponding data have been retained in the analysis presented in the manuscript.

**Supplementary Table 2.** Quantitative data on all proteins identified with an FDR below 1% for each brain region. Columns A-E contain protein identification information. Columns G-R contain the Protein ID for each protein in each regions from our Bayesian analysis, and the N number for each protein from the raw ProteinPilot analysis for each of the six regions. Columns T-Y list the number of distinct peptides associated with each protein in each region, and columns AA-AF contains the number of distinct MS/MS spectra associated with each protein for the purposes of quantification. The remaining column contain the quantitative data for each region, as described in Supplementary Information and the description of our online database.

**Supplementary Table 3.** Data used for the construction of the Edwards-Venn diagram in Figure 2. All genes identified in all six regions are listed in column A. Columns D-I show our call in terms of differential expression between AD and control. Columns K-P show a filtered list of genes whose expression changes in each region. Columns S-U list each changing gene according to the intersection in the Edwards-Venn diagram to which they were assigned.

**Supplementary Table 4.** Output from pathway-enrichment analysis. For each brain region detailed data regarding the pathway enrichment analysis is presented. For each region, significant pathways

(P<0.05) are presented and are ordered into related groups. For each pathway the p-value is presented along with the percentage of protein in that patway which are shown to be differentially expressed and a IPS-generated z-score which is designed to calculate the relative activation (or inhibition) of each pathway, where possible. All genes associated with that pathway are listed and colour-coded according to whether they are upregulated in AD (red) or down-regulated (green). The total numbers of up and down-regulated proteins in each pathway are also provided.

**Supplementary Table 5.** Results from DAVID analysis of pathways enriched specifically in Cerebellum.
