## Supplementary figures and images for "Regional protein expression in human Alzheimer’s brain correlates with disease severity"

### Supplementary Materials

Xu et al. - Supplementary Figure 1

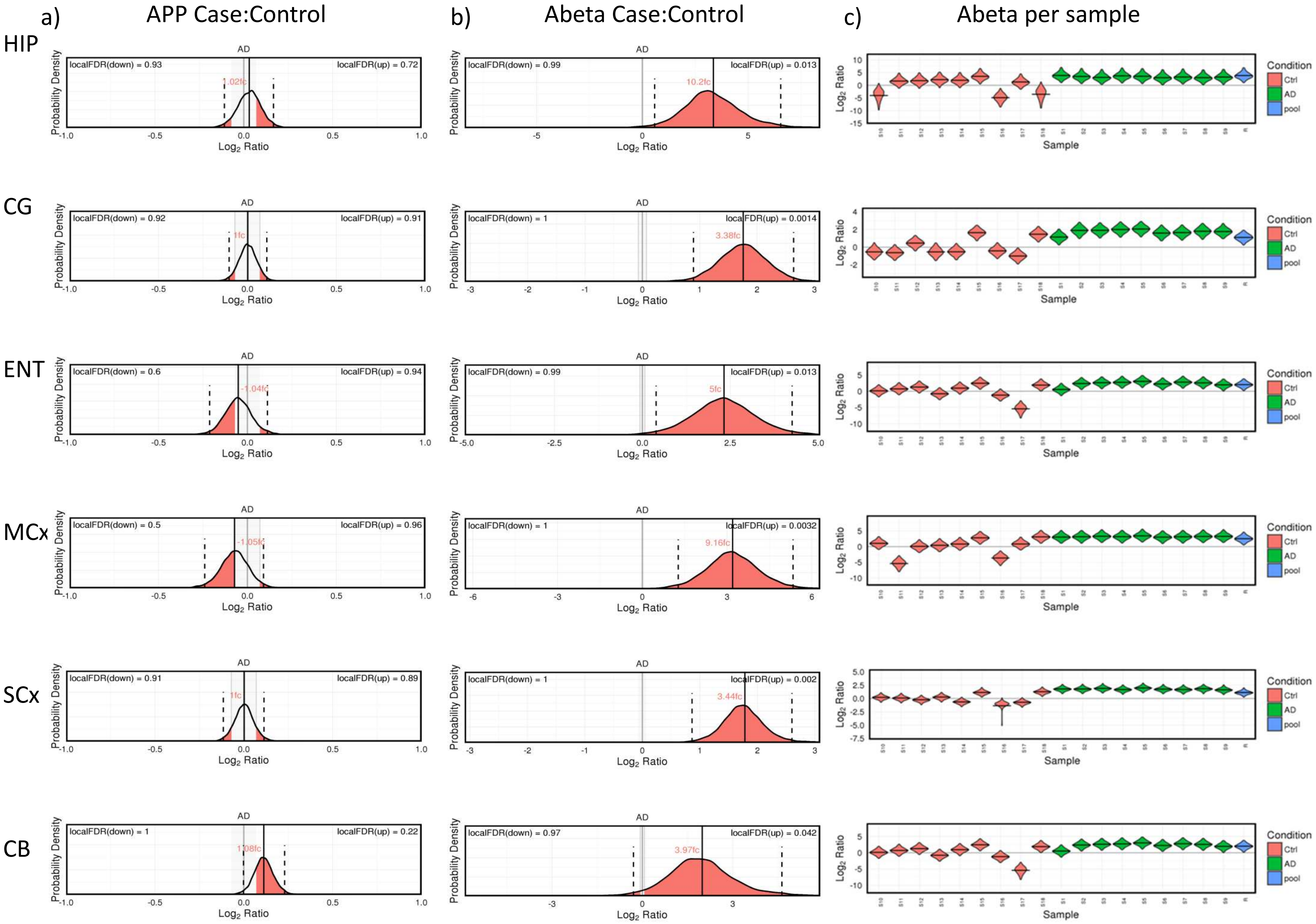
